# A viral genome packaging motor transitions between cyclic and helical symmetry to translocate dsDNA

**DOI:** 10.1101/2020.05.23.112524

**Authors:** Michael Woodson, Joshua Pajak, Wei Zhao, Wei Zhang, Gaurav Arya, Mark A. White, Paul. J. Jardine, Marc C. Morais

## Abstract

Molecular segregation and biopolymer manipulation require the action of molecular motors to do work by applying directional forces to macromolecules. The additional strand conserved E (ASCE) ring motors are an ancient family of molecular motors responsible for diverse tasks ranging from biological polymer manipulation (e.g. protein degradation and chromosome segregation) to establishing and maintaining proton gradients across mitochondrial membranes. Viruses also utilize ASCE segregation motors to package their genomes into their protein capsids and serve as accessible experimental systems due to their relative simplicity. We show by CryoEM focused image reconstruction that ASCE ATPases in viral dsDNA packaging motors adopt helical symmetry complementary to their dsDNA substrates. Together with previous data, including structural results showing these ATPases in planar ring conformations, our results suggest that these motors cycle between helical and planar cyclical symmetry, providing a possible mechanism for directional translocation of DNA. We further note that similar changes in quaternary structure have been observed for proteasome and helicase motors, suggesting an ancient and common mechanism of force generation that has been adapted for specific tasks over the course of evolution.

## INTRODUCTION

A fundamental step in the life cycle of a virus is packaging the viral genome within a protective protein capsid. In one encapsidation strategy, genome compaction and capsid assembly occur simultaneously as shell proteins assemble around the condensing genome. This strategy is employed by most enveloped RNA viruses such as alpha viruses and flaviruses (*1*–*3*). In a second strategy, an empty virus capsid is first assembled, and the genome is then actively translocated into this pre-formed container. This strategy is utilized by dsRNA viruses (*4*, *5*), most large ssDNA viruses (*6*) and virtually all dsDNA viruses such as herpes virus, pox virus, adenovirus, and all the tailed dsDNA bacteriophages (*7*, *8*). This second enapsidation strategy requires that nascent viruses overcome substantial enthalpic and entropic barriers resisting DNA compaction to condense DNA within the confined space of the capsid. Thus, dsDNA viruses have evolved specialized molecular machinery to efficiently package their genomes (*7*–*9*).

The molecular motors that power genome encapsidation are some of the most powerful motors in nature, capable of producing forces in excess of 50 piconewtons (*10*–*13*). For reference, myosin and kinesin each operate at around 5 to 10 pN (*14*–*16*), the approximate magnitude of force required to break a hydrogen bond. The high forces generated by dsDNA packaging motors are necessary to work against the ~20 atm of pressure permeating capsids at the later stages of encapsidation (*17*). Energy for packaging is provided by a virus-encoded ATPase that converts chemical energy released by ATP binding and hydrolysis into mechanical translocation of DNA (*8*, *9*). These ATPases reside on the additional-strand catalytic glutamate (ASCE) branch of the ancient and ubiquitous P-loop NTPase superfamily (*18*–*20*). Often, arranged as homomeric rings, ASCE ATPases are typically involved in polymer manipulation tasks (e.g. protein degradation, chromosome segregation, DNA recombination, DNA strand separation, and conjugation), or in molecular segregation (e.g. proton movement by the F1-ATP synthase) (*21*). Hence, insights into the mechanism of viral DNA packaging will also illuminate the mechanistic principles of a broad class of molecular motors responsible for basic macromolecular partitioning processes.

Bacteriophage phi29 has emerged as a powerful experimental system for investigating genome packaging motors, and development of an efficient *in vitro* packaging system in phi29 has facilitated interrogation via multiple experimental approaches (*7*, *22*, *23*). Genetic, biochemical, and structural results show that the motor is comprised of three components (Figure 1A) (*7*, *24*): 1) a dodecameric portal, or connector protein (gene product 10 (gp10))(*25*); **2)** a pentameric ring of a phage encoded structural RNA molecule (pRNA) (*25*–*29*); and **3)** a pentameric P-loop ASCE ATPase ring motor (gene product 16 (gp16)), analogous to the large terminases in other phage systems that drive packaging (*24*, *25*, *30*, *31*). The components assemble as three co-axial rings at a unique vertex of the capsid, and the dsDNA genome is threaded through a continuous channel along their common central axis and into the viral capsid (Figure 1A).

**Figure 1.**
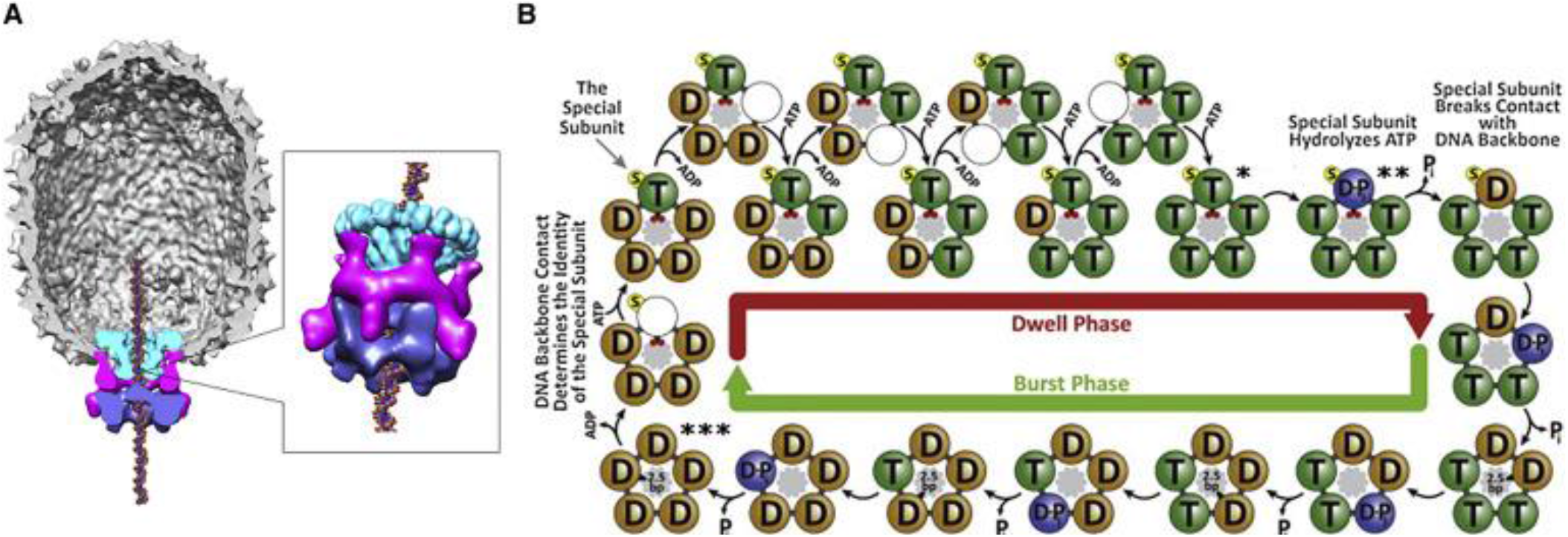
The Bacteriophage phi29 dsDNA Packaging Motor: **(A)** Cut-away side view of the bacteriophage phi29 dsDNA packaging motor as determined by cryoEM. Molecular envelopes of the connector, pRNA, and ATPaseare shown in green, magenta, and blue, respectively. **(B)** Model of the mechano-chemical cycle of the packaging motor as determined by single-molecule experiments (adapted from Chistol et al., 2012). The top and bottom halves of the panel show the chemical and mechanical transitions in the dwell phase and burst phases, respectively. Selected points of the mechano-chemical cycle that are specifically referred to in the text are marked by asterisks.

Biochemical and single molecule analysis indicate that the packaging motor operates in a highly coordinated manner (*12*, *17*, *32*, *33*) (Figure 1B). The mechano-chemical cycle is separated into two distinct phases (Figure 1B) (*33*). During the burst phase, four gp16 subunits sequentially hydrolyze ATP, resulting in translocation of 10 bp of DNA in four 2.5 bp steps (*33*). The role of the fifth subunit is poorly understood, but it has been proposed to play a regulatory role in aligning the motor with DNA for the next translocation burst ((*17*, *34*). No DNA translocation occurs during the subsequent dwell phase, when all five gp16 subunits release ADP from the previous hydrolysis cycle and load ATP in an interlaced manner. More recent optical laser tweezer experiments show that nucleotide exchange during the dwell is coordinated between adjacent subunits, and that the order of exchange is influenced by subunit interactions with DNA (*35*). Using altered DNA substrates, additional single molecule experiments indicate that the motor makes two types of contact with the translocating DNA (*36*). During the dwell, the motor makes electrostatic contacts with the phosphate backbone of the DNA, whereas the motor was proposed to use a ‘steric paddle’ to make non-specific contacts during the burst phase to actively translocate the DNA. It has also been shown that DNA rotates 14° during each 10 bp translocation burst, likely to maintain motor/substrate alignment. Finally, it has been shown that both the magnitude of this rotation and the step-size of the motor change in response to head filling at the last stages of packaging (*17*). While extensive biochemical and single molecule analysis have provided a detailed kinetic scheme describing what happens during packaging (Figure 1B), the physical basis of force generation and subunit coordination remain largely unknown. Visualizing the molecular events that constitute the mechano-chemical cycle requires structural characterization of the motor at distinct points along this pathway. Toward this end, the structure of phi29 particles stalled during DNA packaging by the addition of the non-hydrolyzable substrate γ-S-ATP was determined by cryo-electron microscopy. While the overall resolution of the reconstruction was better than 3 Å, density corresponding to the inherently dynamic motor was somewhat worse. Nonetheless, focused reconstruction and local averaging of density corresponding to various motor components resulted in local resolutions ranging from ~3.8 Å for the portal to ~7 Å for the ATPase component of the motor. At these resolutions, it was possible to unambiguously fit atomic resolution structures of the connector, pRNA, and ATPase into the map and to refine these structures against their corresponding densities. The resulting structure is the first complete near-atomic resolution structure of a viral dsDNA packaging motor and shows that the N-terminal domain is arranged as a cracked ring/one-turn helix that tracks the helical symmetry of dsDNA. Together with previous structural, biochemical, biophysical, and single molecule data, these results suggest a molecular basis for DNA translocation.

## RESULTS

Presently, it is only possible to assemble functional phi29 DNA packaging motors in the context of the entire procapsid (*7*, *22*, *23*). While the structure of a prohead-motor complex could theoretically be determined by X-ray crystallography, it has not yet been possible to crystallize such a complex for any virus or phage. Further, the ~14 MDa mass of this complex is well beyond the reach of NMR. While cryoEM has emerged as a powerful technique to obtain near atomic resolution structures of large complexes (*37*, *38*), it is still challenging to resolve inherently flexible components (e.g. molecular motors) to high resolution. This challenge is more formidable for dsDNA packaging motors attached to virus procapsids since there is a symmetry mismatch between the relatively small motor components and the massive icosahedral capsid on which they reside. Reconstructing such an unbalanced symmetry mismatch is still non-trivial (*4*, *26*, *29*, *39*–*41*). Hence, due to its size and complexity, the most tractable path to the atomic structure of a functional viral dsDNA packaging motor is to determine atomic resolution structures of isolated components by X-ray/NMR/SPA-cryoEM, and then fit them into sub-nm cryoEM reconstructions of the entire motor complex assembled on the procapsid (*42*). To date, we have determined atomic resolution structures of nearly every component of the phi29 packaging motor including the portal (*25*), the prohead-binding domain of the pRNA (*27*), the CCA bulge (*43*) and A-helix of the pRNA (unpublished, Ailong Ke), the N-terminal ATPase domain of gp16 (*31*), and, most recently, the C-terminal vestigial nuclease domain of gp16 (Mahler et al., manuscript in preparation). Further, we have recently determining the full-length structure of a homologous packaging ATPase from a relative of phi29, bacteriophage asccphi28 (Morais lab, unpublished data).

### CryoEM reconstruction of particles stalled during DNA packaging

Having completed the library of atomic resolution structures of phi29 motor components, we turned to single particle cryoEM to image functional motors assembled on procapsids (Figure 2A). Briefly, we incubated 120-base pRNA proheads with gp16, phi29 genomic DNA, ATP, and magnesium for approximately 5 minutes before stopping the packaging reaction via addition of an excess of the non-hydrolysable ATP analog γ-S-ATP. This incubation period is long enough to allow gp16 to assemble on proheads and for functional motors to fully package their genomic DNA substrates. The addition of γ-S-ATP serves two purposes: **1)** it stabilizes packaged particles such that the motor remains attached and the DNA is retained in heads; and **2)** it drives the motor towards a uniform, substrate bound state. Presumably, addition of γ-S-ATP arrests the motors near the end of the dwell phase, when all five subunits are bound to substrate (or non-hydrolysable substrate analog γ-S-ATP) and the motor is poised to enter the burst (Figure 1B, double asterisk).

**Figure 2.**
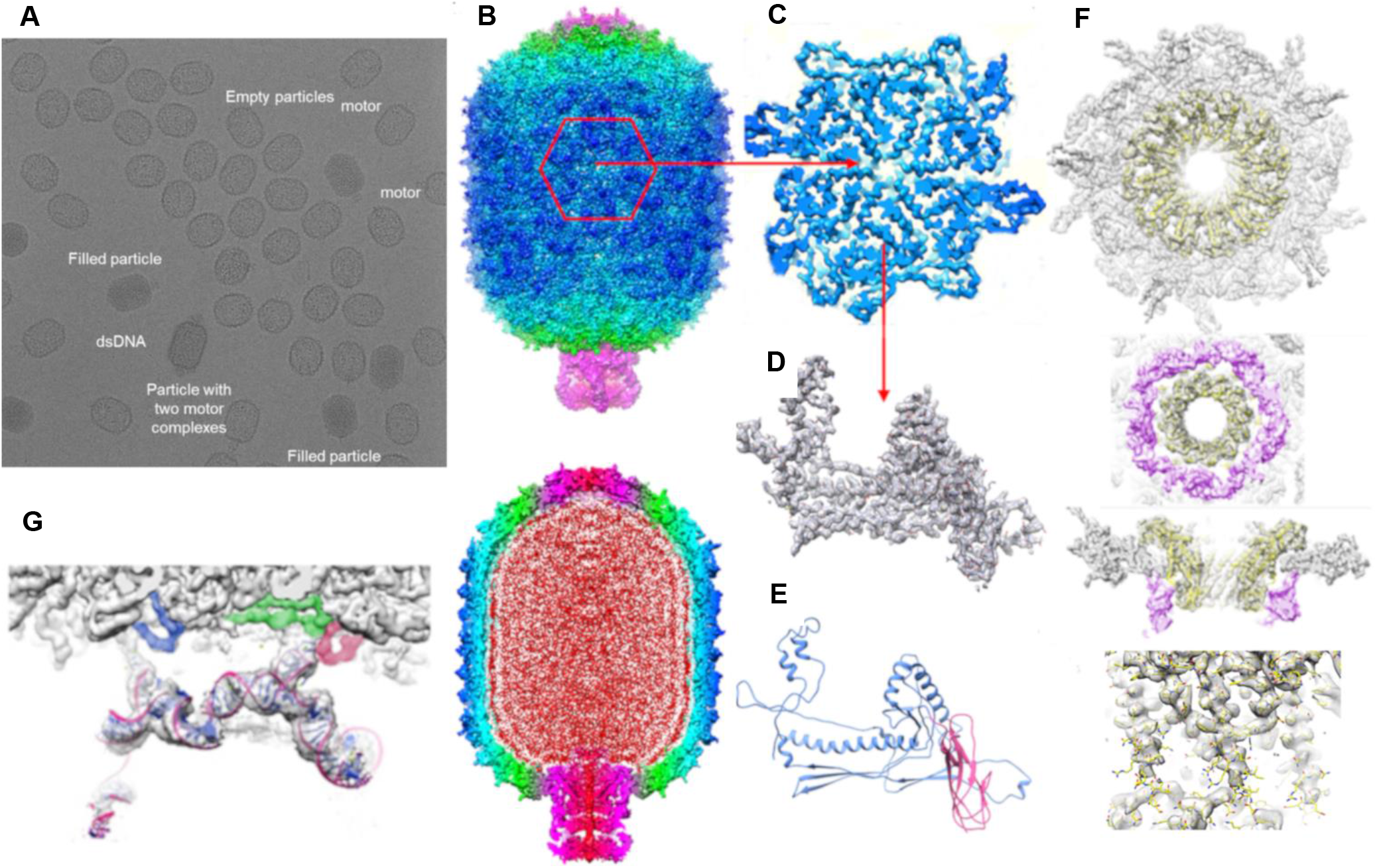
CryoEM reconstruction of bacteriophage phi29 particles stalled during DNA packaging: **A)** CryoEM micrograph of phi29 particles stalled during packaging via addition of γ-S-ATP. Empty and filled particles, unpackaged dsDNA, packaging motors at the unique vertex of procapsids, including an aberrant particle with two packaging motors attached, are indicated. **(B)** Side (top) and cut-away side (bottom) views of particles. Density for the shell is colored by cylindrical radius, changing from red at low cylindrical radii to blue at higher cylindrical radii. The components of the DNA packaging motor at the bottom of particles in and are colored in magenta. Layers of packaged DNA are colored red in the lower panel. **(C)** One hexamer in the T=3, Q=5 prolate icosahedral lattice, outlined in red in (B), is shown at a larger scale. **(D)** Density for one monomer in the hexamer shown in (C) is extracted and shown with the atomic model of the phi29 capsid protein built into its corresponding density. **(E)** Ribbon diagram of the phi29 capsid protein with the N-terminal HK97 domain shown in blue and the C-terminal IG-like domain shown in pink. **(F)** Focused reconstruction of the phi29 portal vertex with the atomic model of the portal built into its corresponding density. The top two panels show end-on views looking from inside the capsid (upper panel), from below the portal (upper middle), side view showing the portal, pRNA, and capsid, and close side view of the central helical region of the portal. **(G)** Atomic resolution structure of the pRNA (RNA backbone shown in magenta and nucleotide bases shown in blue) built into its corresponding density. The E-loop of the pRNA attaches to E-loops in the Capsid protein, clored blue, green and red.

### Procapsid structure

Particles were initially reconstructed assuming C5 symmetry. The resulting map was of high quality (Figure 2B, C), as evidenced by readily recognizable secondary structures and often clear and unambiguous sidechain density (Figure 2D). The overall resolution of the map is ~2.9 Å as estimated by “gold standard” Fourier Shell correlation (FSC) between independently processed half-data sets (*44*, *45*). Consistent with previous reconstructions of phi29 particles (*46*, *47*), the capsid appears to have pseudo-D5 symmetry that is broken by the unique motor vertex at one end of the otherwise T=3 Q=5 prolate icosahedral shell. Density for the capsid was particularly good, and nearly the entire length of the capsid protein density could be traced, and sidechains unambiguously positioned (Figure 2D, E). Further details of model building and refinement of the procapsid will be described elsewhere, but Figure 2B-E shows the overall quality of the density corresponding to the protein shell.

### Symmetry breaking, portal structure

As would be expected, density for the connector was initially poor. Due to the symmetry mismatch between the dodecameric connector and the 5-fold symmetric vertex where it sits, imposing C5 symmetry causes the portal to be incoherently averaged around its 12-fold symmetry axis (*29*). Hence, to improve density for the portal protein, we used a variation of focused reconstruction and symmetry breaking (*29*, *39*, *41*, *48*). Briefly, density corresponding to connector was first flattened by setting corresponding voxel values to the average background value. The resulting map, which no longer has density corresponding to the portal, was projected in the known orientation of each particle that was included in the reconstruction and then translated according to the three rotational and two translational parameters that define each particle’s orientation and center. The density height of each particle projection was scaled to the density height of its corresponding experimental image, and then subtracted from that image. The resulting images should thus only have density corresponding to the portal. Areas corresponding to the portal were extracted using a smaller box and processed as a new data set in Relion (*49*) assuming C12 symmetry.

To reconstruct the portal in the context of the other motor components and the procapsid, we reanalyzed the original images using a variation of a standard symmetry breaking procedures. Since the 12-fold axis of the portal is coincident with the 5-fold axis of the procapsid, the orientations of the whole particle should differ from orientations of extracted portals only in one of the three angles that define particle orientation. Further, this angle can be determined by simply comparing 12 symmetry related projections around the C12/C5 axis to the experimental data. Using this approach, we were able to determine orientations for each particle where both the procapsid and portal were aligned for inter-particle averaging during 3D reconstruction. Density for both the portal and the capsid were excellent in the resulting reconstruction (~3.9 Å resolution; Figure 2G), but detailed results, including model building and interpretations, will be discussed elsewhere. It is not certain why the more traditional symmetry breaking approach didn’t work, wherein 5 symmetry equivalent projections around the C5 axis are compared to experimental images aligned via the procapsid (*29*). However, we suspect that the signal from portal density was insufficient to break symmetry in this case, whereas the massive signal from capsid density could easily break the symmetry when initial orientations were based on the portal.

### pRNA structure

Density corresponding to the pRNA in the initial C5 map was not as clear as density corresponding to the capsid protein but was still easily recognizable as RNA (Figure 2G). Major and minor grooves were apparent with dimensions like those predicted for A-form nucleic acid helices, and serrated periodicity consistent with the phosphate backbone was visible for much of the pRNA. Density for the 74-bases corresponding to the prohead binding domain of the pRNA was better than density corresponding to the A-helix, likely due to increased flexibility of the latter. Density corresponding to the pRNA in the C1 map obtained via symmetry breaking based on portal density described above was similar to density in the original C5 map.

### Symmetry breaking, pRNA-ATPase structure

Since symmetry breaking based on focused reconstruction of the portal resulted in a high-quality reconstruction of the portal itself (Figure 3F) it was surprising that the density corresponding to the ATPase and DNA in this reconstruction was still rather poor (data not shown). The DNA was mostly rod-like, and the only hint of periodicity appeared to be purely translational rather than helical. Density for the ATPase was similarly poor and did not seem much improved relative to a previously published 12 Å map (*31*). A possible explanation is that the motor is highly dynamic, and thus different individual motors are in different conformations and orientations relative to the portal, resulting in smeared density upon inter-particle averaging during 3D reconstruction. Another possibility is that there is yet another symmetry mismatch between the procapsid-portal complex and the ATPase/DNA; even once the portal and procapsid are simultaneously aligned, there are still five different ways that the ATPase and DNA can be arranged with respect to the prohead-portal.

**Figure 3.**
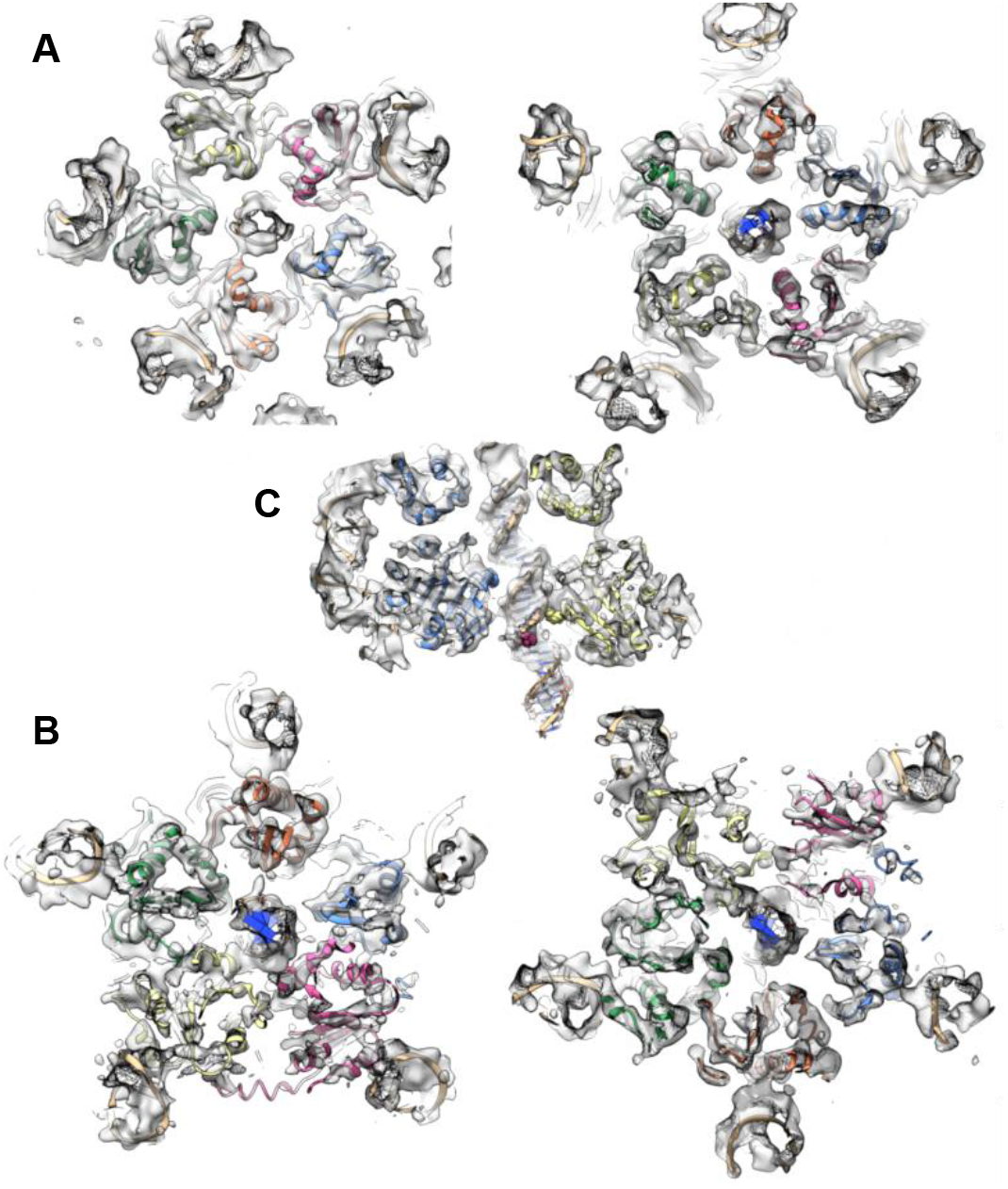
CryoEM reconstruction of the phi29 DNA packaging ATPase motor: **(A)** Density for the CTD planar ring viewed looking from below (left) and above (right). Five copies of the CTD NMR structure (PDB: 6V1W) are fitted into their corresponding densities and shown as red, yellow, green, orange, and blue ribbon diagrams. **(B)** Density for the NTD helical assembly viewed looking from below (left) and above (right). Five copies of the NTD crystal structure (PDB: 5HD9) are fitted into their corresponding densities and shown as red, yellow, green, orange, and blue ribbon diagrams. **(C)** A cut-away side-view of the phi29 ATPase motor, with CTD and NTD ribbon diagrams colored as above. The pRNA (periphery) and DNA (center) are shown as tan ribbon diagrams in all three panels.

To test this possibility, we repeated the symmetry breaking procedure described above except that instead of focusing exclusively on the portal, we focused on density that includes the portal, the pRNA, the DNA, and the ATPase. Further, since DNA in the capsid is not well resolved, it is impossible to correctly subtract DNA density from experimental images. Hence, we excluded views where the capsid is projected onto the motor, and only included particles where the ATPase was clearly visible. Since these confounding views were eliminated and the symmetry breaking mass now included all four motor components, we suspected their combined mass would provide enough signal to differentiate between 5 pseudo symmetry-equivalent positions as in typical symmetry breaking. Indeed, this was the case, as the density for the pRNA, DNA, and ATPase were greatly improved in the resulting map (Figure 3). While the resulting resolution was not high enough to identify side chains or trace the polypeptide chain *ab initio*, it was sufficient to clearly see secondary structural elements in the ATPase and the major and minor grooves in the DNA and pRNA. The phosphate backbone was still readily apparent in the pRNA density and was now visible in parts of the DNA density as well. The overall resolution of the map of the motor vertex was estimated to be ~ 7 Å by FSC (*44*, *45*) and D99 criterion (*50*). Interestingly, density for the portal was not nearly as well resolved as density for the pRNA, DNA, and ATPase (data not shown). While density in the radial direction was well resolved, density in circumferential directions was rather poor, indicating that particles were incoherently averaged around the 12-fold symmetry axis of the portal. This result suggests that symmetry breaking was dominated by signal from the pRNA, ATPase, and DNA, and that the portal density was not aligned in the selected orientations. This could occur if there is no defined relationship between the portal and the motor. Alternatively, there could be five different ways to arrange the portal with respect to the ATPase, but the small mass of the portal could not provide enough signal to successfully classify these different arrangements. We suspect the latter to be true.

### Building an atomic model of the motor

While the separate PDBs for the prohead binding and A-helix (unpublished) domains could be fit into their corresponding densities, these structures had engineered mutations to facilitate crystallization. Hence, since density for the pRNA was of reasonably high quality, a complete 120-base pRNA model was built directly into its corresponding density. However, it proved useful to utilize base-pairing patterns observed in the crystallographic structures as restraints when refining the model against density. Similarly, an atomic model of the DNA was built by first fitting coordinates of a stretch of ideal B-form DNA into its density. Further, the improved density allowed us to unambiguously fit our crystal structure of the N-terminal ATPase domain (NTD: PDB 5HD9) and our NMR structure of the C-terminal vestigial nuclease domain (CTD: PDB 6V1W) into their corresponding densities (Figure 3). The linker region between the domains was built directly into density based on the equivalent linker region in our recently determined full-length ATPase structure of a homolog of gp16, the dsDNA packaging ATPase from bacteriophage asccphi28 (Morais lab, unpublished). Fitted structures were refined against their corresponding densities using the real space cryoEM refinement module in Phenix (*50*). After initial refinement, the map was improved via local averaging over equivalent sub-volumes corresponding to the N- and C-terminal domains of the ATPase, and to the prohead-binding and A-helix domains of the pRNA. The refined models were then manually adjusted to fit this “locally averaged” map, and then refined together against the original, unaveraged map, as in the application of NCS averaging in X-ray crystallography.

### Gross features of the packaging motor

Several gross features of the motor structure are immediately apparent upon an examination of the refined coordinates (Figure 4). First, the pRNA and ATPase are pentameric assemblies, definitively resolving a long-standing controversy regarding the symmetry of these components and refuting several reports of hexameric assemblies (*7*) (Figure 6C). Second, consistent with previous results (*31*), the CTD sits above the NTD and just below the portal where it is wedged between the pRNA and the incoming DNA (Figure 4B). This arrangement of ATPase domains differs from an arrangement reported for the T4 DNA packaging motor, where structural data suggested a reversed domain polarity wherein the NTD attaches to the portal with the CTD hanging just below (*51*). Also, both the N- and C-terminal domains appear to interact with incoming DNA (Figures 3, 4), again differing from results for T4 that suggest only the CTD interacts with DNA (*51*). Additionally, all five NTDs and CTDs appear to interact with DNA, in contrast to previous results we reported that only one subunit contacts the DNA (*35*). Further, the DNA structure differs considerably from canonical B-form DNA. It is stretched/partially unwound in some places, compressed in others, and has a prominent kink between the CTD and NTD, deflecting the direction of the DNA ~ 15° from the expected direction of DNA translocation (Figure 3, 4B, C). Hence, mechanisms requiring strict helical symmetry of B-form DNA may need to be re-evaluated.

**Figure 4.**
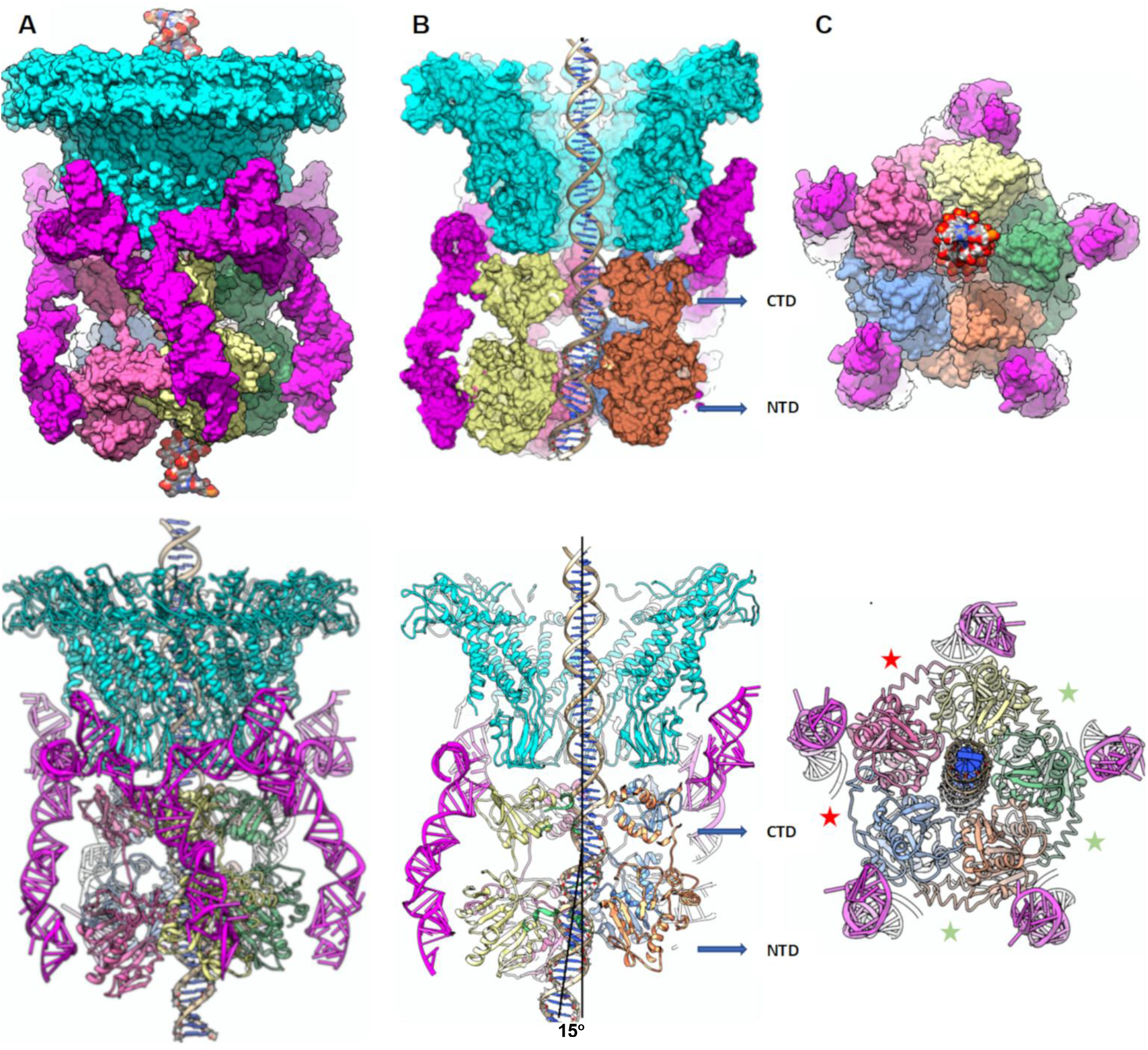
Structure of the bacteriophage phi29 dsDNA packaging motor stalled during packaging: Structure of the packaging motor rendered as molecular surfaces (top panels) and ribbon diagrams (bottom panels) shown from: **(A)** side-view; **(B)** Cut-away side view to visualize the DNA in the central channel; and **(C)** an end-on view, looking from below the motor. The portal and pRNA are colored cyan and magenta, respectively, and the five ATPase subunits are colored yellow, red, blue, orange, and green. Approximate levels of the C-terminal (CTD) and N-terminal (NTD) domains are indicated by blue arrows in panel. Deviations from the rotational component of helical symmetry are shown indicating loose and tight interfaces by red and green asterisks, respectively in (C); the loose interfaces are on either side of the lowest subunit in the ATPase helix, and correspond to the two active sites where there is no clear denisity for ATP. Two black lines in the bottom half of panel (B) are drawn approximately coincident with the helical axis of the DNA before and after the kink that occurs between the CTD and NTD; the ~12.5° angle between the lines is indicated.

While the pRNA and CTD have approximate C5 symmetry, the NTD of the ATPase is not arranged as a planar ring, but rather adopts an arrangement more akin to a cracked ring or a short, one-turn helical assembly (Figures 4, 5, S1, and https://www.youtube.com/watch?v=D2wZOoSJIng). The five NTDs in the ‘helix’ wrap roughly one turn (~10 bp) of dsDNA and are thus arranged such that the two helices are approximately in register approximately every ~2 base pairs. Further, the NTD does not adopt a perfect helical arrangement, as there is some variation in both the twist and rise relating adjacent subunits (Figures 4,5), with subunits S1 and S5 at the top and bottom of the helix differing most in regard to both twist and rise. Since the NTDs are identical, it is not entirely clear how these deviations arise. A possible partial explanation is that the helical arrangement of NTDs results from subunit interactions with DNA, and thus the observed distortions in pitch, twist and direction of the DNA give rise to complementary deviations in the NTD helix. This would explain why S5 deviates most; it sits at the bottom of the helix where the DNA is furthest offset. However, it does not explain why S1 deviates since it is positioned at the top of the helix where the DNA offset is minimal. However, it neighbors S5, and so the deviation of S5 could influence the position of S1. Additionally, while all subunit NTDs contact the pRNA via a small N-terminal helix, S1’s NTD makes an additional contact with the pRNA via a small loop (Figure 5E). This second anchor point may cause S1’s position to deviate from helical symmetry.

**Figure 5.**
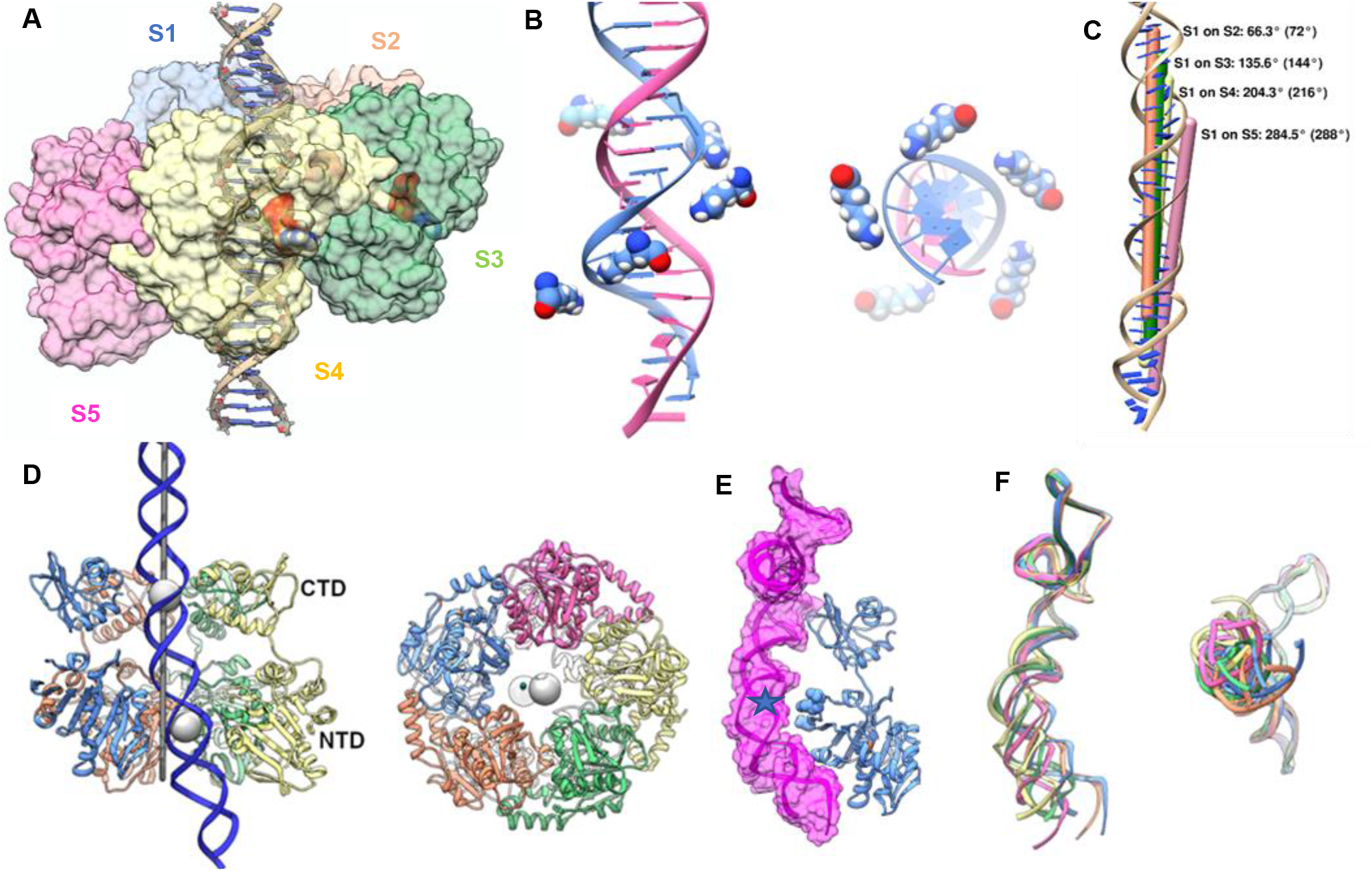
Helical arrangement of ATPase N-terminal domains, offset of NTD and CTD rings, and K56-DNA interactions: **(A)** Side-view of the ATPase with CTDs removed to emphasize the helical arrangement of NTDs; the procapsid, and thus direction of DNA translocation is towards the top of the page. The NTDs from five different subunits are colored differently and shown as semi-translucent surfaces. ATP is also rendered as a molecular surface, colored by element; **(B)** Lysine K56 is arranged as a spiral that approximately tracks the DNA helix. The bottom four subunits from S2, S3, S4 and S5 in panel A track the 3’-5’ strand approximately every 2 bps. Due to the imperfect helical symmetry of the ATPase, the top K56 from S1 is closer to the complementary 3’-5’ strand. **(C)** Helical symmetry axes for superposition of subunits S1 on subunits S2, S3, S4, and S5 are shown as different colored rods and labeled along with their actual and ideal (parentheses) rotations. The translational components of the helical operations are represented by different offsets of the rods, **(D)** Centers of mass (COM) of the NTD and CTD are shown as grey spheres from side and end on views. The 5-fold axis of the phage is shown as a darker grey rod, Subunits are colored as above. **(E)** Subunit S1 (blue ribbon) and its associated pRNA (magenta ribbon in magenta translucent surface). The unique NTD-pRNA contact in S1 is shown as a blue star. **(F)** superpositions of the pRNA from side and end on views. The pRNAs were superimposed onto pRNA-S1 via their prohead binding domains (bases 1-74) and colored according to their associated NTDs. Note that the blue pRNA-S1 is bent such that its distal NTD-binding end regions moves both up and to the right.

Since both the CTD and pRNA are arranged as planar rings, one question that arises is how the NTD can assume a helical arrangement when its anchor points are planar. While each NTD could bind to a different section of the pRNA to give rise to a staggered structure, our results suggest otherwise. Ignoring the unique 3^rd^ contact with pRNA made by S1, all subunits essentially bind pRNA the same way. Instead, the pRNA changes conformation to adapt to the helical arrangement of NTDs (Figure 5F). In this way the pRNA functions analogously to the suspension of an off-road vehicle, where shock absorbers compress, extend, and flex to allow the wheels of the vehicle to maintain contact with the ground over uneven terrain. Similarly, the pRNA allows the NTD to maintain contact with its helical DNA substrate. However, unlike off-roader’s shock absorber, the A-form RNA helices cannot extend or compress, and thus positional adjustments of the NTD are limited to bending and flexing motions. Thus, to ‘raise’ the position of S1 to the top of the helix, the pRNA bends rather than simply shortening. As a result, in addition to positioning S1 closer to the portal, it is also slightly displaced in a direction ~perpendicular to the 5-fold axis of the phage. Since S1 is part of the larger NTD assembly, the whole cracked ring should be displaced in this same direction. The results presented here are consistent with this scenario. The pRNA attached to S1 is the most bent (Figure 5E), as it must be to maintain S1’s position at the top of the helix. Further, plotting the centers of mass (COM) of the NTD and CTD show that while the CTD COM lies on the global 5-fold axis, the NTD COM is displaced as predicted (Figure 5D). This explains why the DNA is bent; since the DNA is nestled in the central pore of the NTD cracked ring, it is also displaced when the NTD cracked ring is displaced as it adopts a helical structure. Hence, there is likely a complex interplay between the NTD adopting a helical structure to bind DNA and the deformation of DNA structure in response to this binding.

### Interactions with DNA

The central pores of both the CTD ring and the NTD helix of the ATPase are rich in positively charged and polar residues that are well-positioned to interact with DNA (Figures 3–5). Given the resolution of the map, it is difficult to definitively catalog these residues, but using a distance cutoff of 4 Å to select residues that could potentially interact with DNA shows that while the motor interacts with both strands, there seem to be more interactions with one of the two strands. This observation is consistent with single molecule results showing that the motor tracks along the 5’-3’ strand during translocation (*36*). The overall helical character of the NTD described above is particularly apparent if one focuses on a single residue in the motor pore. Figure 5B shows K56, which appears to track along the 5’-3’ strand of the dsDNA helix approximately every 2 to 2.5 base pairs. Note that due to deviations from helical symmetry in both the dsDNA and the NTD cracked ring/helix, each K56 interacts with the DNA in a slightly different way. This is especially true for S1 at the top of the helix, which seems to be better positioned to interact with the opposite 3’-5’ strand, consistent with its deviation from strict C5 helical symmetry discussed above.

### Inter-molecular interactions between motor components

The N- and C-terminal domains are connected by a long linker analogous to the “lid” domains in other ASCE ATPases (Figure 6). In addition to connecting the N- and C-terminal domains, the NTD-CTD linker also ‘links’ adjacent subunits. It folds into a small three-helix domain, where the middle helix interacts with an adjacent subunit. Because of deviations in the helical pitch and twist, the orientation and the distance spanned by the linker arm is not uniform around the helix. Distance and orientation of the linker seems to be adjusted by varying the length of the two helices flanking the helix that binds the neighboring subunit; more overall helical content shortens the distances, whereas direction can be altered by ending the helical region at different extents around a helical turn (Figures 4C, 6C). The density for the linker regions from the subunits at the top (S1) and bottom (S5) of the ATPase helix is poor compared to the other three linker arms. This may be due to increased dynamics and/or conformational heterogeneity of these two linkers due to their unique positions in the NTD helix. In particular, the linker originating from the top subunit S1 must reach down to interact with the bottom subunit S5, and therefore must adopt a significantly different orientation compared to the other subunits. Subunit S5 at the bottom of the helix is also in a unique position and is part of the split interface between subunits S1 and S5. However, S5’s linker is not part of this interface, but instead is part of the interface between subunits S5 and S4. Hence, the environment of S5’s linker should be like the environments of the three preceding subunits S4, S3, and S2. Nonetheless, like the top subunit, the bottom subunit’s overall position and orientation deviate from the stricter symmetry observed for the inner three subunits (Figures 4C, 5C, 6C). This difference in position and orientation may account for part of the increased flexibility observed for this region (see also below).

**Figure 6.**
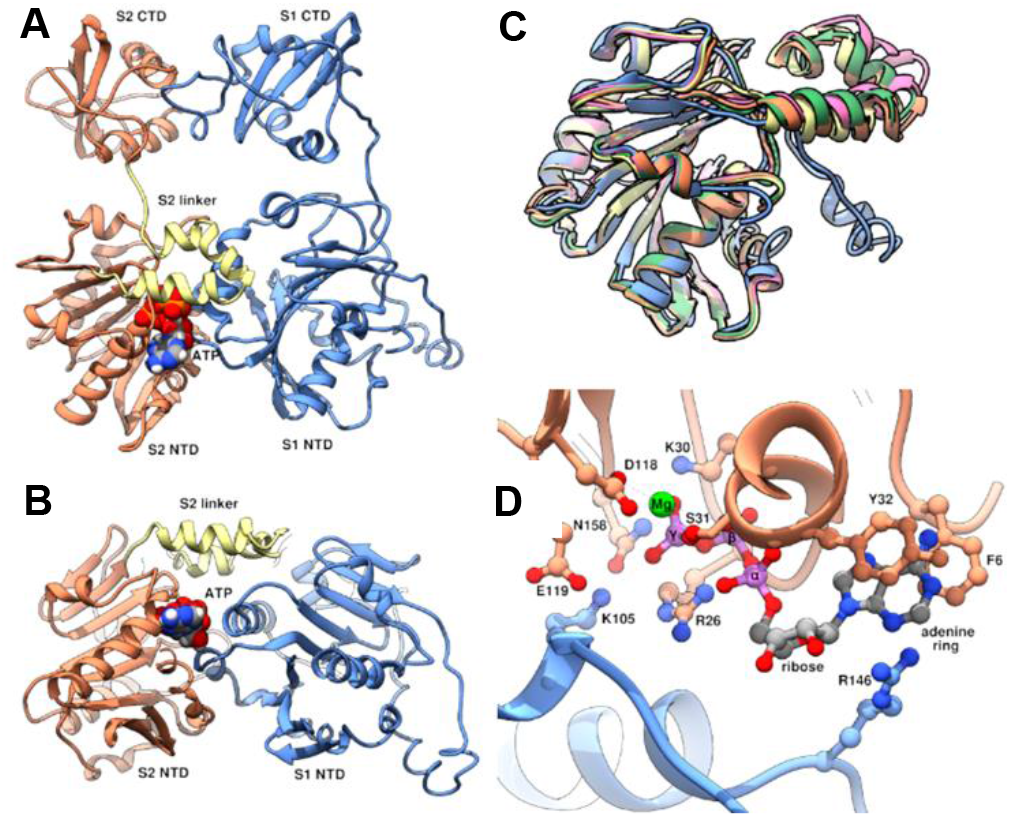
Domain linker and trans-acting residues: **(A)** Side-view and **(B)** end-on view of adjacent subunits S1 and S2, colored as in figures 6 and 7. NTDs and CTDs are labeled, and the linker domain in S2 is highlighted in yellow. ATP is shown as space filling spheres colored by element **(C)** Superposition of the NTDs of all five subunits to illustrate structural variability of the linker domain. Note that the relative orientation of linker domains from subunits S1 (blue), and S5 (pink) differ the most, and correspond to subunits at the top (S1) and bottom (S5) of the ATPase helix. **(D)** Close up of the active site between subunits S1 and S2. Residues important for binding and/or catalysis, including K105 and R146, are shown as ball and stick figures, colored by heteroatom, and labeled. ATP atoms are shown as ball and stick, colored by element; note phosphorous is colored light purple rather than the typical orange to facilitate visibility.

### Active site of the ATPase

Although the resolution of the motor-vertex map is not sufficient to definitively visualize ATP, nucleotide binding sites of ASCE ATPases are well characterized and the location of ATP can be inferred by superimposing the structure of a related ATPase complexed with ATP. In this case, superimposing the structure of the bacteriophage P74-26 DNA packaging ATPase complexed with substrate analog ADP-BeF_3_ (*52*) allowed positioning of substrate in active sites located between two adjacent subunits, consistent with previous data (*31*). In the middle three subunits (S2, S3, S4), the nucleotide fits well into otherwise unexplainable density. However, in the top and bottom subunits (S1 and S5), there is no significant density corresponding to the expected position of the nucleotide. . Additionally, the linker arm closes over the active site, and is thus positioned to respond to ATP-binding, hydrolysis, and product release (Figures 6A,B). Density for the linker arm in S1 and S5 is poor, suggesting that nucleotide binding might influence the position and flexibility of the linker.

As described above, the active site resides between two subunits in the NTD helix. Thus, residues from two adjacent subunits contribute to a single active site and each contribute to ATP binding and hydrolysis (Figure 6D). In the ‘cis’-acting subunit, ATP is positioned near the Walker A and B motifs that reside on loops connecting strands on one edge of the central beta sheet of the ASCE fold, consistent with previous results reported for phi29 and other ATPases. ATP is further stabilized by positively charged residues contributed in ‘trans’ by an adjacent subunit. In particular, the ATPs seem to be sandwiched between R146 and K105 (Figure 6D). In ring ATPases, an arginine residue often acts in trans to trigger ATP hydrolysis either by stabilizing the negative charge that accumulates on the phosphate oxygens during the hydrolytic transition state or by similarly stabilizing the negative charge on the ADP leaving group. Based on sequence alignment with members of the FtsK branch of the ASCE superfamily, R146 was predicted to be the canonical ATPase ‘arginine finger’. This prediction was supported by biochemical experiments designed to report trans-acting activity (*31*, *35*). While the current structural data could support R146 acting as the arginine finger, K105 seems equally well, if not better, positioned to assume this role. Specifically, K105 is positioned nearer the γ-phosphate of ATP (where nucleophilic attack occurs), whereas R146 is closer to the adenine base and pentose sugar (Figure 6D). Further, an arginine residue equivalent to K105 in the thermophilic bacteriophage P74-26 was shown to function as the arginine finger (*52*). Hence, while R146 likely does act in trans during nucleotide cycling, assigning it the catalytic-trigger role of the arginine finger may be premature. Indeed, single molecule optical laser tweezer experiments suggest the R146 may play a role in nucleotide exchange (*35*), thus explaining the reported trans-activity while allowing for the possibility that K105 could function as the catalytic ‘arginine’ finger in gp16. Of note, in addition to closing over the active site, the linker interacts with the edge of the sheet in the trans-acting subunit where R146 and K105 reside (Figures 6B, D), consistent with potential roles in coordinating either/both ATP hydrolysis and nucleotide exchange in adjacent subunits.

## DISCUSSION

### Structural comparison of the phi29 packaging motor and other ASCE ATPases

While it is well known that all ASCE ATPases share a modified Rossman fold and are thus similar at the tertiary level, the extent of similarity at the quaternary level is less clear. Early crystallographic and EM structures of ASCE ATPases without their polymeric substrates bound indicated that most were assembled as planar hexameric rings. More recently, several structures of ASCE ATPases complexed with their polymeric substrates have been determined to near-atomic resolution by single particle cryoEM (*53*). Many of these structures have a helical arrangement of subunits rather than a planar-ring organization. Based on these structures, a hand-over-hand mechanism has been proposed for polymer translocation, wherein nucleotide hydrolysis is coupled to the movement of a subunit from one end of the helix to the other such that the ATPase helix slithers along it polymer substrate in a hand-over-hand, spiral-escalator type mechanism (*54*–*57*).

### Mechanism of translocation

Despite assembling as a pentamer rather than a hexamer, the results presented here show that like other ASCE ATPases, the NTDs of the phi29 DNA packaging ATPase adopt a helical structure when engaged with their polymeric dsDNA substrate. Such an arrangement suggests that phi29 might also employ a hand-over-hand, spiral-escalator-type mechanism to propel its dsDNA into the procapsid. While there are many attractive aspects of such a mechanism for viral dsDNA packaging motors, there are also some concerns. Primarily, the well-characterized dwell-burst behavior observed for phi29 in single molecule experiments does not naturally emerge from such a mechanism. Escalators, whether linear or helical, are continuous, and do not pause as the last stairstep reaches the top/bottom. Further, like the hexameric ASCE ATPases, the phi29 ATPase has also been observed in planar configurations (*24*, *30*). Similarly, our recently determined X-ray and cryoEM-SPA structures of the full-length packaging ATPase from the phi29 relative ascc-phi28 in three different nucleotide states (apo, ADP-, AMPPNP-bound) are all planar rings in absence of DNA (unpublished data).

We propose an alternative mechanism for DNA packaging based on the helical NTD structure observed here and planar structures described above. The essence of the mechanism is that the NTD of the ATPase cycles between helical and planar symmetry, and that DNA is driven into the procapsid as the subunits adopt the planar configuration (Figure 7, S2: https://www.youtube.com/watch?v=v3k35VYTuFI&feature=youtu.be). For simplicity’s sake, deviations from strict helical symmetry in the NTD, distortions of the DNA structure and direction, and offset of the NTD relative to the CTD were not considered, and positions of NTD subunits have been adjusted to obey strict helical/planar C5 symmetry in figure 7 and the accompanying linked video.

**Figure 7.**
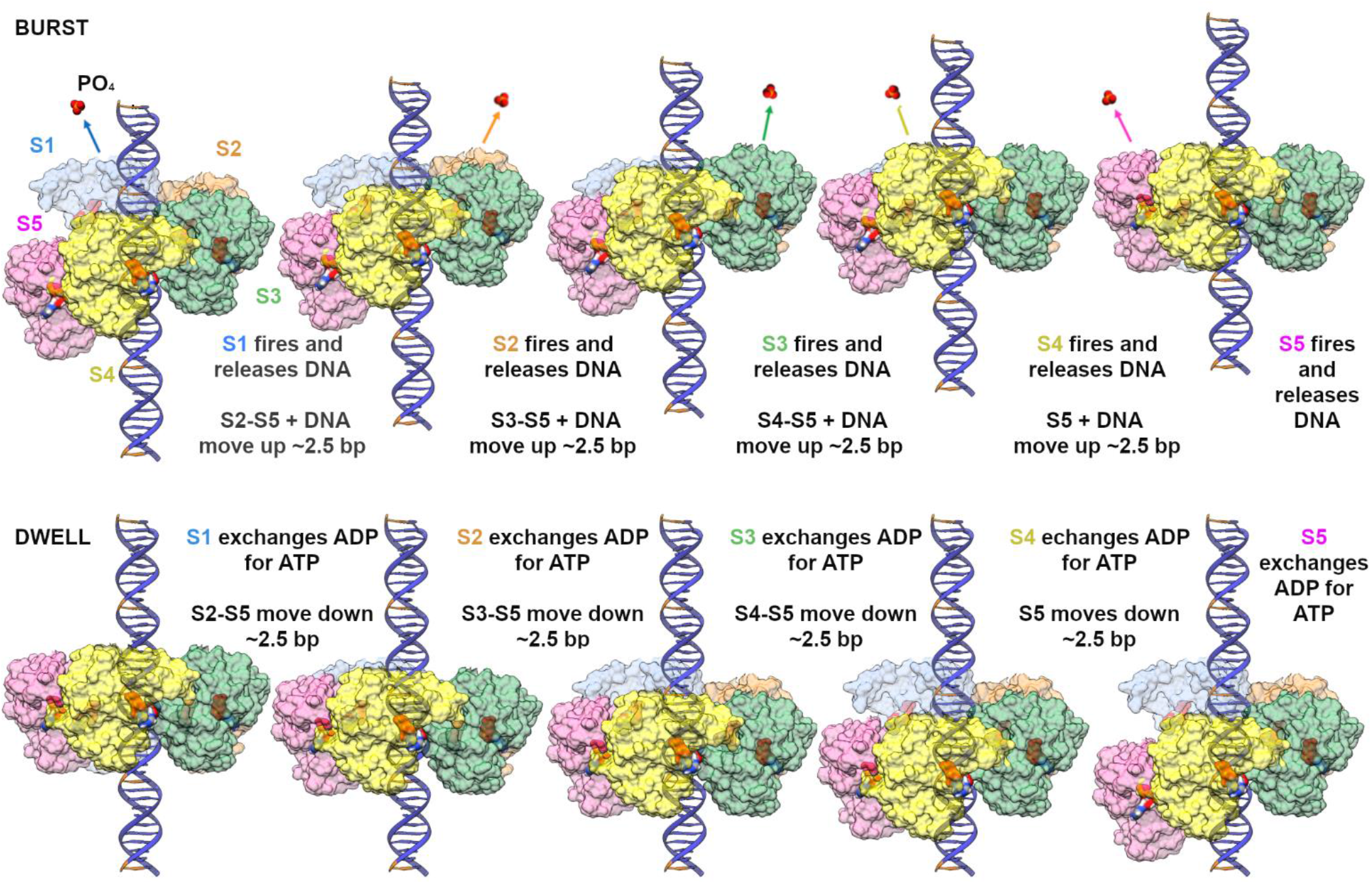
Mechanism of DNA translocation: For simplicity, only the NTDs of the ATPase are shown, and subunit positions have been adjusted to obey perfect helical symmetry as described in the text. NTDs from the five different subunits, labeled S1-S5, are colored differently and shown as semi-translucent surfaces. ATP and PO_4_ are rendered as opaque molecular surfaces, colored by element. DNA is shown in blue, with the B-form DNA helical repeat indicated by coloring every 10 bps on the 5’-3’ strand orange. The top panel shows the burst phase; ATP hydrolysis causes the NTDs to transition from a helical to a planar configuration and drive DNA into the procapsid (top of page). Note that while the first four hydrolysis events move 10 bp of DNA in four 2.5 bp steps, the last hydrolysis event in S5 does <u>not</u> translocate DNA. The bottom panel illustrates the dwell, when sequential exchange of ADP for ATP occurs and the motor resets to the helical configuration as NTD subunits S2-S5 walk down DNA. Note that during the dwell, there is no translocation of DNA.

The mechano-chemical cycle begins when all five ATPase subunits have bound ATP at the end of the dwell. As with many ASCE ATPases, gp16 has a high affinity for DNA when ATP is bound and lower affinities in ADP-bound and apo states (*53*, *58*). To optimally engage its DNA substrate, the NTD adopts a helical arrangement that is approximately complementary to the DNA. In the first step of the burst, S1 at the top of the helix hydrolyzes ATP, releasing its grip on DNA and causing subunits S2-S5 to move a distance ~equal to the rise of the NTD helix (~7 to 8.5 Å), thus bringing S2 into the same plane as S1. Further, since subunits S2-S5 have not yet hydrolyzed ATP, they remain tightly bound to DNA, causing DNA to also translocate ~2 to 2.5 bp into the procapsid. Once S2 is in plane with S1, trans-acting K105 in S1 is positioned to trigger hydrolysis in S2. As S2 hydrolyzes ATP, it lets go of DNA and the S3-S5:DNA complex now moves as a unit. This results in translocation of another ~ 2 to 2.5 bp of DNA and places S3 in the plane of the ring such that hydrolysis in S3 can be triggered in-trans by K105 from S2. This process continues to permute around the ring until the burst ends as S4 fires, completing the helix-to-planar transition and resulting in translocation of ~1 helical turn of dsDNA (~10 bp). At this point, S5 has yet to fire. Since the helix-to-planar transition is complete, firing at S5 will not translocate DNA any further; all subunits are in plane with S1, and thus there is no impetus for any to move higher. However, S4’s trans-acting K105 will nonetheless be positioned to trigger hydrolysis in S5. Further, since the motor reset requires that subunits S2-S5 return to their previous positions along the NTD helix, S5 likely needs to hydrolyze ATP to release its grip on DNA. Otherwise DNA would be pulled out of the capsid as S5 migrates back to the bottom of the NTD helix. Hence, at the end of the burst/beginning of the dwell, S5 likely hydrolyzes ATP, allowing it to release DNA.

Single molecule experiments have shown that nucleotide exchange occurs sequentially and is highly coordinated, with one subunit completing exchange of ADP for ATP prior to exchange in the next subunit (*33*, *35*) (Figure 1B). The order of nucleotide exchange around the ring could occur in either the same or opposite direction as hydrolysis. While we cannot rule out either path, we favor exchange occurring in the same direction (Figure 7, bottom). In this path, as S5 comes into plane with S1 at the end of the burst, its trans-acting R146 would be positioned to initiate nucleotide exchange in S1. Further, since DNA has moved one helical repeat, S1 is positioned to re-bind DNA upon completion of exchange. Exchange in S1 would then promote exchange in S2, and so on until each subunit has reassumed its previous position in the NTD helix. A sensible aspect of this direction of exchange is that when S2 and subsequent subunits sequentially drop down one position in the NTD helix, they would be optimally aligned to bind to DNA as they re-bind ATP and increase their affinity for DNA. In contrast, if exchange occurred in the opposite direction and S5 were to initiate the walk down the helix, S5 would become competent to bind DNA before it completes its downward journey. Thus, this path could result in S5 pulling DNA backwards out of the capsid as the motor resets the helical NTD arrangement. However, it is likely that subunits prefer a particular orientation of the DNA backbone relative to the NTD. If so, then this optimal relative arrangement would not occur until S5 had returned to the bottom of the helix. Indeed, in this direction of exchange, S2 through S5 would all reach the optimal binding position at the same time, resulting in a zipper-like interaction between the NTD helix and DNA, possibly providing a signal that the dwell has ended.

A few gross features of this mechanism are worth noting. First, S1 acts as a pivot point for the helix-to-planar transition, and thus maintains its position at the top of the helix throughout the mechano-chemical cycle. Additionally, ATP hydrolysis in one subunit does not impose a force directly on DNA, but rather translocates DNA by causing movement of adjacent DNA-bound subunits. This motion is likely actuated by changing the length and orientation of the moving subunit’s linker arm, as described above. Regardless, only subunit S5 maintains contact with the DNA throughout the entire burst phase. A similar mechanism was proposed based on single molecule results (*33*) that predicted the ATPase moved DNA by transitioning between a planar and cracked ring while one subunit, equivalent to S5, held onto the DNA.

The proposed mechanism provides explanations for some previously puzzling data. For example, the well-characterized dwell-burst cycle (*33*) emerges naturally from a helix-to-planar mechanism; no translocation occurs during nucleotide exchange as the motor walks back down the NTD helix. Additionally, this mechanism explains how a five-subunit motor evolved to utilize a 4-stroke cycle; only 4 steps are required to convert a 5-subunit helix into a planar ring. Similarly, a previously proposed “special” subunit is consistent with the role of either subunit S1, which is unique in its translationally invariant position at the top of the helix, or S5, wherein hydrolysis of ATP is not coupled to DNA translocation. Additionally, the step size of the motor is set by the rise of the NTD helix, and thus need not be coupled to an integral number of base pairs.

Even apparent disagreements between single molecule results and our structural results may be copacetic. Single molecule results show that each burst is composed of four ~8.5 Å steps (*33*). However, the translational components of the step wise transformations to superimpose adjacent subunits along the NTD helix (and thus the motor step sizes), range from ~6 to ~7.5 Å. A possible explanation for this small discrepancy is that the NTD helix tries to adapt itself to the symmetry of its polymeric substrate, and thus the shorter steps may result from DNA distortions, including DNA bending, deviations from ideal B-form DNA, and the radial offset of the NTD cracked ring. Indeed, despite these distortions, subunits S2-S5 interact with DNA approximately every 2 bp along the 3’-5’ strand, which is consistent with single molecule results (*36*). On a related note, DNA was held under tension in the single molecule experiments, which may affect its ability to bend and/or distort as observed here, resulting in a slightly different NTD helical arrangement. Additionally, our structure represents the state of the motor at the end of packaging, where single molecule results have demonstrated that the mean step size of the motor decreases in response to head-filling (*17*). Hence the supposed discrepancies may not be as significant as they seem.

The proposed mechanism also makes several testable predictions. First, we would predict that a structure of the motor stalled with ADP would show that the NTD is in a planar ring configuration. Second, the mechanism predicts that the motor could also package dsRNA, but that the magnitude of the burst would reflect the periodicity of A-form RNA. Further, we would predict that a structure of particles stalled while packaging dsRNA with a substrate analog would show that the NTD adopts a helical arrangement more complementary to an A-form RNA helix.

One question that arises for any proposed dwell-burst mechanism is what prevents the DNA from sliding out of the head during the dwell. Assuming that we are indeed imaging molecules at the end of the dwell, one possibility is that the observed contacts between the CTD and DNA act as a valve to prevent DNA slippage. A second possibility is related to the observed deviations from strict helical symmetry, and the resulting NTD-DNA contacts. While subunits S2-S5 interact with the 5’-3’ strand approximately every two base pairs at the end of the dwell/beginning or the burst, S1 is better positioned to interact with the opposite strand. According to the mechanism outlined above, S5 remains attached to the DNA for the duration of the translocation burst. Further, since S1 has not moved and DNA has translocated one helical turn, S1 will again be positioned to interact with the DNA as it was at the beginning of the burst. In this way, once S1 exchanges ADP for ATP, it can bind the DNA again and hold it in place while the motor resets. Thus, the observed deviation from helical symmetry would allow S1 to interact with DNA in both the helical and planar configurations, providing a means of holding DNA during the dwell.

Additionally, the details of how hydrolysis in one subunit causes the neighboring subunit (and its bound DNA) to move up are not yet clear. However, the calculated buried surface area between two adjacent subunits in the planar ring arrangement is greater than that for two adjacent subunits in the helical arrangement. Since binding energy is roughly proportional to buried surface area, this provides a thermodynamic motive for the transition. Extending this idea over planar-ring and cracked ring/helical symmetries suggests an energetic framework for understanding the motor’s mechano-chemical cycle. Consider the energy well for two proteins binding in an optimal way. If more than two subunits can bind this way in a higher symmetry structure, the energy wells for each binding pair add in phase, resulting in deep energy well for the assembly. In the mechanism proposed here, the motor transitions from helical to planar symmetry during the burst, and then from planar back to helical symmetry during the dwell. Having a motor that transitions between symmetric structures ensures that once a burst or dwell is initiated, it goes to completion to occupy a deep energy well associated with a symmetric structure.

However, the motor must not become trapped at the end of the burst or dwell. Thus, the motor uses ATP binding and hydrolysis, coupled with DNA binding, to both climb out of energy wells and to change their relative depths. As mentioned above, there is more buried surface area in the planar ring assembly than in the helical assembly. Not only is there more buried surface area per interface, there are more interfaces since considerably less surface area is buried at the split interface in the cracked ring. Thus, in the absence of ATP and DNA, the planar ring is more energetically favorable. In the presence of substrate DNA, the energy for each subunit is the sum of the energy of interaction with its neighbors plus the energy of its interaction with DNA. In the ADP-bound or apo state, the affinity for DNA is low, and thus DNA interactions contribute little – the planar ring remains lower energy. However, when ATP is bound, affinity for DNA is high enough that the energy gained via DNA interaction is more than enough to offset the difference in energy between planar and helical interfaces. Hence, during the dwell, as ATP replaces ADP, the additional energy gained by DNA binding causes the helical structure to be the lower energy well, and thus the subunits adopt the helical/cracked ring conformation. Upon ATP hydrolysis during the burst, there is little affinity for DNA, and the planar ring structure becomes the deeper energy well again. Thus, as ATP is hydrolyzed, released, and re-bound, the relative depths of the energy wells associated with helical and planar arrangements shift, and the motor traverses the oscillating energetic landscape much like a child’s slinky toy walks down a set of stairs. These general principles may be applicable to other systems, including other ring ATPases that engage polymeric structures where planar and helical/cracked ring structures have been observed (*54*–*57*).

## METHODS

### Production of Packaging Components

*Bacillus subtilis* 12A (*sup*-) host cells were infected with the phi29 mutant phage *sus* 8.5(900)-16(300)-14(1241), which is defective in the packaging ATPase and thus cannot package DNA. The resulting empty prohead particles were purified via ultra-centrifugation using sucrose gradients as previously described (*59*, *60*). To ensure that the pRNA associated with these particles was structurally and compositionally homogeneous, purified particles were first treated with RNase A to remove any attached pRNA and re-purified as previously described (*60*). In parallel, 120b pRNA was produced from the plasmid pRT71 by *in vitro* transcription using T7 RNA polymerase and purified by denaturing urea polyacrylamide gel electrophoresis as previously described (*61*). The RNA-free proheads were then reconstituted with uniform 120b as previously described (*60*). The genomic DNA-gp3 substrate and packaging ATPase gp16 were prepared and purified as previously described (*31*, *59*).

### DNA packaging assay

The *in vitro* DNA packaging assay is based on a DNase protection assay and was performed as described previously (*59*, *60*). Briefly, proheads (8.3 nM), DNA-gp3 (4.2 nM), and either wild-type or mutant gp16 (166 to 208 nM) were mixed together in 0.5X TMS buffer in 20 μl and incubated for 5 min at room temperature. ATP was then added to 0.5 mM to initiate packaging. After 15 min incubation, the mixture was treated with DNase I (1 μg/ml final) and incubated for 10 min to digest the unpackaged DNA. An EDTA/Proteinase K mix was then added to the reaction mixture (25mM and 500 μg/ml final concentration, respectively) and incubated for 30 min at 65°C to inactivate the DNase I and release the protected, packaged DNA from particles. The packaged DNA was analyzed by agarose gel electrophoresis. Packaging efficiency was calculated by densitometry using a UVP Gel Documentation System.

### ATPase assays

ATPase activity of prohead/gp16 (wt or mutant) motor complexes was determined by measuring production of inorganic phosphate by the EnzChek Phosphate Assay Kit (Life Technologies) as described previously (*43*). Briefly, a reaction mixture containing reaction buffer (either kit buffer [50mM Tris, pH 7.5, 1mM MgCl_2_, 0.1mM sodium azide] or TM buffer [25 mM Tris, pH7.6, 5 mM MgCl_2_]), 0.2 mM of MESG (2-amino-6-mercapto-7-methylpurine riboside) with proheads (4.2nM) and gp16 (125nM) in 90μl was preincubated at room temperature for 10 min in the presence of PNP (purine nucleoside phosphorylase, 0.1 unit). ATP was added to 1mM to initiate the reaction and production of Pi measured in the spectrophotometer at 360 absorbance for 10 min. For the complementation assay, the D/E and R146A mutants were mixed in the ratios indicated to yield the 125nM gp16 concentration.

### Production of DNA Packaging Intermediates for cryoEM

To generate the stalled DNA packaging intermediates for cryoEM reconstruction, packaging reactions consisting of DNA-gp3, RNA-free proheads reconstituted with 120b pRNA and gp16 were assembled at 2X concentration (see above) and ATP was added to initiate DNA packaging. Three minutes post-ATP addition, gamma-S-ATP was added to 100μM (Roche) (ATP concentration is 500μM) and incubated for 2 minutes. To maximize packaging efficiency, a prohead-to-DNA ratio of 2:1 was used. Hence, at most, only 50% of the particles could package DNA. However, far fewer particles had actually packaged DNA (Figure 2), and thus there was considerable amounts of unpackaged DNA fibers in the background of cryoEM images. Thus, to facilitate freezing and remove background noise contributed by unpackaged DNA, 1 unit of RQ DNase I (Promega) was added and incubated at room temperature for 10 minutes. The sample was placed on ice until grid preparation for cryoEM imaging.

While enough gp16 was added to assemble functional motors on every particle, the large number of unpackaged particles raised concerns that motors on these particles were either not fully (or properly) assembled. Thus, the reaction mixture was loaded on a sucrose gradient (with an excess of gamma-S-ATP included) to separate DNA-filled particles from empty particles. The gradient band corresponding to filled particles was dialyzed against TMS buffer supplemented with gamma-S-ATP to remove sucrose, concentrated, and loaded onto cryoEM grids for freezing and imaging. The resulting grids had a good distribution of almost entirely filled particles. Since most particles in the images had packaged DNA, we could presume that their packaging motors were functional. Although a small amount of “unpackaged” DNA was still present, this likely resulted from some particles losing their DNA during purification and was not enough to hinder image processing. Although these empty particles had likely packaged but then lost DNA, they were nonetheless excluded from image processing to ensure only functional motors in the same state were reconstructed.

### CryoEM grid preparation, data collection, image processing and model building

Approximately 3 μl of prohead particles stalled during packaging (see above production of packaging intermediates for cryoEM) was applied to quantifoil 2X2 holey carbon grids prior to plunge freezing. Sample grids were flash-frozen in liquid ethane cooled to liquid nitrogen temperatures on holey carbon grids using a vitrobot automated virification system from Thermo Fisher. Data were collected on the Titan Krios microscope housed at the Electron Imaging Center for Nanomachines (EICN) at UCLA. Data were collected as previously described (*31*). Individual movie frames were aligned with Motioncor2 (*62*), and particles were picked with EMAN2 (*63*). Subsequent image processing steps, including symmetry breaking and focused reconstruction, were carried using Relion (*49*) and/or symmetry breaking scripts written for Relion (*39*). Atomic models were fitted and/or manually built into their corresponding densities using COOT (*64*) and refined using PHENIX (*65*). All maps and PDBs are available upon request and will be released to the public via the PDB and EMDB upon acceptance in a peer reviewed journal.

## ACKNOWLEDGMENTS

This work was supported by Public Health Service grant GM122979 (to P.J.J. and M.C.M.), GM127365 (to M.C.M.), and GM11817 (to G.A.). CryoEM data collection was supported by U24GM116 (Midwest CryoEM Consortium) from the National Institutes of Health. We would also like to acknowledge the Sealy Center for Structural Biology and Molecular Biophysics (SCSB) for support of the UTMB cryoEM and computational core facilities, and the Electron Imaging Center for Nanomachines (EICN) at UCLA for use of the Titan Krios microscope.

## REFERENCES

1. S. Apte-Sengupta, D. Sirohi, R. J. Kuhn, Coupling of replication and assembly in flaviviruses. Curr Opin Virol 9, 134–142 (2014).

2. A. Mendes, R. J. Kuhn, Alphavirus Nucleocapsid Packaging and Assembly. Viruses 10, (2018).

3. S. Sotcheff, A. Routh, Understanding Flavivirus Capsid Protein Functions: The Tip of the Iceberg. Pathogens 9, (2020).

4. S. L. Ilca et al., Multiple liquid crystalline geometries of highly compacted nucleic acid in a dsRNA virus. Nature 570, 252–256 (2019).

5. E. J. Mancini, R. Tuma, Mechanism of RNA packaging motor. Advances in experimental medicine and biology 726, 609–629 (2012).

6. L. L. Ilang et al., DNA packaging intermediates of bacteriophage phi X174. Structure 3, 353–363 (1995).

7. M. C. Morais, The dsDNA packaging motor in bacteriophage o29. Advances in experimental medicine and biology 726, 511–547 (2012).

8. V. B. Rao, M. Feiss, Mechanisms of DNA Packaging by Large Double-Stranded DNA Viruses. Annu Rev Virol 2, 351–378 (2015).

9. V. B. Rao, M. Feiss, The bacteriophage DNA packaging motor. Annu Rev Genet 42, 647–681 (2008).

10. D. N. Fuller, D. M. Raymer, V. I. Kottadiel, V. B. Rao, D. E. Smith, Single phage T4 DNA packaging motors exhibit large force generation, high velocity, and dynamic variability. Proc Natl Acad Sci U S A 104, 16868–16873 (2007).

11. D. N. Fuller et al., Measurements of single DNA molecule packaging dynamics in bacteriophage lambda reveal high forces, high motor processivity, and capsid transformations. J Mol Biol 373, 1113–1122 (2007).

12. S. Liu, G. Chistol, C. Bustamante, Mechanical operation and intersubunit coordination of ring-shaped molecular motors: insights from single-molecule studies. Biophys J 106, 1844–1858 (2014).

13. D. E. Smith et al., The bacteriophage straight phi29 portal motor can package DNA against a large internal force. Nature 413, 748–752 (2001).

14. J. T. Finer, R. M. Simmons, J. A. Spudich, Single myosin molecule mechanics: piconewton forces and nanometre steps. Nature 368, 113–119 (1994).

15. E. Meyhofer, J. Howard, The force generated by a single kinesin molecule against an elastic load. Proc Natl Acad Sci U S A 92, 574–578 (1995).

16. K. Svoboda, C. F. Schmidt, B. J. Schnapp, S. M. Block, Direct observation of kinesin stepping by optical trapping interferometry. Nature 365, 721–727 (1993).

17. S. Liu et al., A viral packaging motor varies its DNA rotation and step size to preserve subunit coordination as the capsid fills. Cell 157, 702–713 (2014).

18. L. M. Iyer, D. D. Leipe, E. V. Koonin, L. Aravind, Evolutionary history and higher order classification of AAA+ ATPases. J Struct Biol 146, 11–31 (2004).

19. L. M. Iyer, K. S. Makarova, E. V. Koonin, L. Aravind, Comparative genomics of the FtsK-HerA superfamily of pumping ATPases: implications for the origins of chromosome segregation, cell division and viral capsid packaging. Nucleic Acids Res 32, 5260–5279 (2004).

20. D. D. Leipe, Y. I. Wolf, E. V. Koonin, L. Aravind, Classification and evolution of P-loop GTPases and related ATPases. J Mol Biol 317, 41–72 (2002).

21. A. M. Burroughs, L. M. Iyer, L. Aravind, Comparative genomics and evolutionary trajectories of viral ATP dependent DNA-packaging systems. Genome Dyn 3, 48–65 (2007).

22. S. Grimes, P. J. Jardine, D. Anderson, Bacteriophage phi 29 DNA packaging. Adv Virus Res 58, 255–294 (2002).

23. P. Guo, S. Grimes, D. Anderson, A defined system for in vitro packaging of DNA-gp3 of the Bacillus subtilis bacteriophage phi 29. Proc Natl Acad Sci U S A 83, 3505–3509 (1986).

24. M. C. Morais et al., Defining molecular and domain boundaries in the bacteriophage phi29 DNA packaging motor. Structure 16, 1267–1274 (2008).

25. A. A. Simpson et al., Structure of the bacteriophage phi29 DNA packaging motor. Nature 408, 745–750 (2000).

26. S. Cao et al., Insights into the structure and assembly of the bacteriophage 29 double-stranded DNA packaging motor. J Virol 88, 3986–3996 (2014).

27. F. Ding et al., Structure and assembly of the essential RNA ring component of a viral DNA packaging motor. Proc Natl Acad Sci U S A, (2011).

28. P. Guo, C. Peterson, D. Anderson, Prohead and DNA-gp3-dependent ATPase activity of the DNA packaging protein gp16 of bacteriophage phi 29. J Mol Biol 197, 229–236 (1987).

29. M. C. Morais et al., Cryoelectron-microscopy image reconstruction of symmetry mismatches in bacteriophage phi29. J Struct Biol 135, 38–46 (2001).

30. J. S. Koti et al., DNA packaging motor assembly intermediate of bacteriophage phi29. J Mol Biol 381, 1114–1132 (2008).

31. H. Mao et al., Structural and Molecular Basis for Coordination in a Viral DNA Packaging Motor. Cell Rep 14, 2017–2029 (2016).

32. G. Chistol et al., High degree of coordination and division of labor among subunits in a homomeric ring ATPase. Cell 151, 1017–1028 (2012).

33. J. R. Moffitt et al., Intersubunit coordination in a homomeric ring ATPase. Nature 457, 446–450 (2009).

34. K. H. Choi, M. C. Morais, D. L. Anderson, M. G. Rossmann, Determinants of bacteriophage phi29 head morphology. Structure 14, 1723–1727 (2006).

35. S. Tafoya et al., Molecular switch-like regulation enables global subunit coordination in a viral ring ATPase. Proc Natl Acad Sci U S A 115, 7961–7966 (2018).

36. K. Aathavan et al., Substrate interactions and promiscuity in a viral DNA packaging motor. Nature 461, 669–673 (2009).

37. W. Kuhlbrandt, Cryo-EM enters a new era. Elife 3, e03678 (2014).

38. W. Kuhlbrandt, Biochemistry. The resolution revolution. Science 343, 1443–1444 (2014).

39. S. L. Ilca et al., Localized reconstruction of subunits from electron cryomicroscopy images of macromolecular complexes. Nat Commun 6, 8843 (2015).

40. M. C. Morais, Breaking the symmetry of a viral capsid. Proc Natl Acad Sci U S A 113, 11390–11392 (2016).

41. R. Stass, S. L. Ilca, J. T. Huiskonen, Beyond structures of highly symmetric purified viral capsids by cryo-EM. Current opinion in structural biology 52, 25–31 (2018).

42. M. G. Rossmann, M. C. Morais, P. G. Leiman, W. Zhang, Combining X-ray crystallography and electron microscopy. Structure 13, 355–362 (2005).

43. E. Harjes et al., Structure of the RNA claw of the DNA packaging motor of bacteriophage Phi29. Nucleic Acids Res 40, 9953–9963 (2012).

44. M. van Heel et al., Single-particle electron cryo-microscopy: towards atomic resolution. Q Rev Biophys 33, 307–369 (2000).

45. M. van Heel, M. Schatz, Fourier shell correlation threshold criteria. J Struct Biol 151, 250–262 (2005).

46. M. C. Morais et al., Conservation of the capsid structure in tailed dsDNA bacteriophages: the pseudoatomic structure of phi29. Mol Cell 18, 149–159 (2005).

47. Y. Tao et al., Assembly of a tailed bacterial virus and its genome release studied in three dimensions. Cell 95, 431–437 (1998).

48. M. C. Morais et al., Bacteriophage phi29 scaffolding protein gp7 before and after prohead assembly. Nat Struct Biol 10, 572–576 (2003).

49. J. Zivanov et al., New tools for automated high-resolution cryo-EM structure determination in RELION-3. Elife 7, (2018).

50. P. V. Afonine et al., New tools for the analysis and validation of cryo-EM maps and atomic models. Acta Crystallogr D Struct Biol 74, 814–840 (2018).

51. S. Sun et al., The structure of the phage T4 DNA packaging motor suggests a mechanism dependent on electrostatic forces. Cell 135, 1251–1262 (2008).

52. B. J. Hilbert et al., Structure and mechanism of the ATPase that powers viral genome packaging. Proc Natl Acad Sci U S A 112, E3792–3799 (2015).

53. S. Zhang, Y. Mao, AAA+ ATPases in Protein Degradation: Structures, Functions and Mechanisms. Biomolecules 10, (2020).

54. A. H. de la Pena, E. A. Goodall, S. N. Gates, G. C. Lander, A. Martin, Substrate-engaged 26S proteasome structures reveal mechanisms for ATP-hydrolysis-driven translocation. Science 362, (2018).

55. E. J. Enemark, L. Joshua-Tor, Mechanism of DNA translocation in a replicative hexameric helicase. Nature 442, 270–275 (2006).

56. H. Han, C. P. Hill, Structure and mechanism of the ESCRT pathway AAA+ ATPase Vps4. Biochem Soc Trans 47, 37–45 (2019).

57. N. Monroe, H. Han, P. S. Shen, W. I. Sundquist, C. P. Hill, Structural basis of protein translocation by the Vps4-Vta1 AAA ATPase. Elife 6, (2017).

58. Y. R. Chemla et al., Mechanism of force generation of a viral DNA packaging motor. Cell 122, 683–692 (2005).

59. S. Grimes, D. Anderson, The bacteriophage phi29 packaging proteins supercoil the DNA ends. J Mol Biol 266, 901–914 (1997).

60. W. Zhao, M. C. Morais, D. L. Anderson, P. J. Jardine, S. Grimes, Role of the CCA bulge of prohead RNA of bacteriophage o29 in DNA packaging. J Mol Biol 383, 520–528 (2008).

61. R. J. Reid, J. W. Bodley, D. Anderson, Characterization of the prohead-pRNA interaction of bacteriophage phi 29. J Biol Chem 269, 5157–5162 (1994).

62. S. Q. Zheng et al., MotionCor2: anisotropic correction of beam-induced motion for improved cryo-electron microscopy. Nat Methods 14, 331–332 (2017).

63. S. J. Ludtke, P. R. Baldwin, W. Chiu, EMAN: semiautomated software for high-resolution single-particle reconstructions. J Struct Biol 128, 82–97 (1999).

64. P. Emsley, B. Lohkamp, W. G. Scott, K. Cowtan, Features and development of Coot. Acta Crystallogr D Biol Crystallogr 66, 486–501 (2010).

65. P. D. Adams et al., PHENIX: a comprehensive Python-based system for macromolecular structure solution. Acta Crystallogr D Biol Crystallogr 66, 213–221 (2010).

